# Aerial imagery and deep learning accurately estimate maize foliar disease severity

**DOI:** 10.64898/2026.06.03.729887

**Authors:** Cole Hammett, Katelyn Rumley, Peter Balint-Kurti, Joseph L. Gage

## Abstract

Southern leaf blight (SLB) is a foliar disease of maize (Zea mays L.) caused by the necrotrophic fungal pathogen Cochliobolus heterostrophus. Genetic resistance is the most effective control method for SLB. Developing disease resistant maize lines requires field trials during which disease phenotypes must be visually assessed. Remote sensing using drones is an emerging technology that can be leveraged for high-throughput phenotyping of disease severity that is otherwise labor-intensive and subjective. This project used a deep learning approach to estimate SLB disease severity of single-row maize plots from drone imagery. Over 26,000 plot-level images produced from flights conducted across three growing seasons were labeled with in-field visual scores taken contemporaneously by expert raters. Variation in environmental conditions contributed to a labeled image dataset that reflects the complexity of agronomic field experiments. We assessed the ability of nine deep learning models from three architectural families to estimate disease severity. The best-performing model, EVA-02-B, achieved strong cross year generalization (R^2^ = 0.697). Error analysis found that performance was more strongly associated with seasonal disease progression and flight-score time offset than with image-level noise. UAV-based deep learning estimated SLB severity with comparable precision to expert raters. This study lays the groundwork for integrating automated phenotypes into genetic studies of disease resistance.

**PLAIN LANGUAGE SUMMARY:** Southern leaf blight (SLB) of maize is a disease that causes yield loss worldwide and developing resistant varieties offers the best hope for controlling the disease. Studying SLB resistance requires plant pathologists to visually score severity in the field, a labor-intensive method that requires expertise. To address these challenges, we asked whether SLB severity scoring could be automated using drone images and artificial intelligence (AI). We trained AI models using three years of image and score data then compared the results to visual scores taken by five plant pathologists. The best performing AI model showed a similar level of consistency to the experts and proved capable of scoring severity despite unpredictable and uncontrollable conditions that affect field imaging experiments such as weeds or shadows. These findings provide a validated method that improves the efficiency of maize disease research, a critical area of study for agricultural sustainability and productivity.

## 1 INTRODUCTION

Maize (*Zea mays L.*) has the highest production by volume of any staple cereal with extremely versatile uses as feed, food, and biofuel (Erenstein et al., 2022). Diseases pose a major threat to maize production by reducing yield, producing toxins that limit grain use, and increasing management costs (Betts et al., 2025). The development of disease resistant maize varieties offers a reliable and cost-effective solution for controlling diseases and lowering production inputs in economically significant plant pathosystems (Nelson et al., 2018; Wisser et al., 2006). Quantitative disease resistance (QDR) is a preferred management strategy because of its durability, non-race-specific effects, and efficacy against necrotrophic pathogens (Poland et al., 2009; Yang et al., 2017a). However, the moderate effect of multiple genetic loci and strong environmental influence complicate genetic dissection and integration of QDR into elite maize hybrids (Nelson et al., 2018; Yang et al., 2017a). Breeding for QDR requires substantial effort toward in-field disease severity assessments, imposing limits to the number of genotypes, locations, and developmental stages that can be observed. Visual scores are imperfect labels that include both biological signal and rater variation, reducing phenotypic precision and affecting downstream estimates of resistance (Poland & Nelson, 2011). These barriers can be addressed through the development of automated in-field methods for maize field trials that offer higher throughput and greater consistency.

Southern leaf blight (SLB) is a foliar disease in maize caused by the necrotrophic fungal pathogen *Cochliobolus heterostrophus*. SLB is infamous in the United States for the 1970-1971 epidemic when U.S. corn yields suffered an estimated 16% loss due to the widespread adoption of a gene that was key to hybrid seed production at the time and, unfortunately, also a source of susceptibility to a novel race of *C. heterostophus* (Balint-Kurti & Pataky, 2024). SLB infection causes the formation of necrotic lesions beginning on the lower leaves that have variable shape, size, and color influenced by genetic backgrounds among inbreds and hybrids (Munkvold & White, 2016). Resistance to SLB is largely quantitative and relies on a variety of mechanisms (Belcher et al., 2012; Chen et al., 2023; Holley, 1989; Kump et al., 2011; Lim & Hooker, 1976; Yang et al., 2017b). While resistant germplasm has significantly reduced yield loss in the United States, SLB continues to be a significant constraint to maize production in tropical and subtropical regions (Munkvold & White, 2016).

Molecular and computational innovations continuously improve the accessibility and affordability of genetic data collection and analysis. However, the implementation of these powerful methodologies in genetic mapping studies and breeding programs is impeded by the quality and quantity of the phenotypic data and availability of suitable mapping populations (Araus & Cairns, 2014; Gage et al., 2018; Wang et al., 2026; Yang et al., 2020). Several studies have successfully identified quantitative trait loci (QTL) associated with SLB resistance using phenotypes derived from in-field visual scores recorded multiple times per growing season (Balint-Kurti et al., 2007; Bian et al., 2014; Carson et al., 2004; Chen et al., 2023; Kump et al., 2011; Li et al., 2018; Martins et al., 2019; Negeri et al., 2011; Yang et al., 2017b; Zwonitzer et al., 2009). This current standard of visual scoring for SLB severity has limited throughput, precision, and accuracy due to the inherent qualities of the in-field severity measurements (Bock et al., 2020). Improving SLB severity phenotyping has the potential to increase the statistical power and reproducibility of genetic mapping studies for QDR and provide the throughput needed by modern breeding programs (Gage et al., 2018; Poland & Nelson, 2011).

Recent advancements in sensor technology and artificial intelligence (AI) enable the development of high-throughput phenotyping (HTP) methods, offering a non-destructive, scalable and automated alternative to traditional assessment with visual scoring (Wang et al., 2026). While HTP of disease severity is achievable with remote sensing, it often necessitates time-consuming image evaluation or extensive validation (Bock et al., 2022; Singh et al., 2016; Wu et al., 2019). An increasingly popular HTP modality is imagery captured from unmanned aerial vehicles (UAVs) mounted with red-green-blue (RGB) cameras. Feature engineering to extract spectral or textural information from UAV images has been widely adopted, providing data that can be directly used by classical machine learning models. However, this approach overlooks key spatial patterns and suffers from environmental interference. Deep learning (DL), a branch of machine learning inspired by biological neural networks, can learn spatial patterns directly and avoid hand-crafted features with models specialized for image analysis such as convolutional neural networks (CNN) and vision transformers (ViT). This compliments the high-quality RGB image data captured from UAV platforms and facilitates straightforward validation of automated disease severity assessment with disease signals observable to the human eye.

Currently, the use of AI in plant pathology has largely been limited by the availability of sizable and accurately labeled datasets (Jeger et al., 2024). Previous studies that used deep learning to assess maize foliar disease from RGB imagery have focused on object detection or pixel segmentation, typically at the leaf level (Craze et al., 2022; DeChant et al., 2017; Green et al., 2012; Haque et al., 2021; Lee et al., 2025; Singh et al., 2022; Stewart et al., 2019; Wiesner-Hanks et al., 2019; Wu et al., 2019; Yin et al., 2022). Other limitations of current methods include low-throughput handheld imaging devices, destructive leaf sampling, lesion annotations, single leaf or plant targets, and uncoupling from field observations. In this study, we built a three-year dataset of over 26,000 plot-level images labeled with in-field visual scores provided by expert raters. We evaluated the capabilities of nine pre-trained supervised deep learning models across three architecture families (CNN, ViT, hybrid) to estimate SLB severity from our dataset with an agronomically relevant cross-year test evaluation to mirror real-world estimation of unlabeled fields in unseen environments. Comparing visual scores from multiple expert raters to imaged-based scores revealed similar performance between the visual and image-based scoring methods. We then characterize sources of error from flight conditions, image noise, and label imperfections to provide insight into model mechanisms and limitations. Our results demonstrate that UAV-based deep learning can reliably estimate SLB severity with accuracy comparable to visual scorings, and lay groundwork for integrating automated phenotypes into genetic studies of QDR.

## 2 METHODS AND MATERIALS

### 2.1 Experimental Design and Field Assessment

#### 2.1.1 Field Design and Inoculation

All data were collected in fields at the Central Crops Research Station (CCRS) in Clayton, North Carolina during the 2023, 2024, and 2025 growing seasons. Two separate fields within the CCRS were planted each year with 2.43 meter (8 foot) single-row plots with 0.97 meter (38 inch) row spacing. All blocks were planted with a single row of border plots and check lines. Fields were inoculated with isolates 2-16Bm and Hm540 inoculum of *Cochliobolus heterostrophus*, the causal agent of SLB (Carson, 1998). The preparation of fungal inocula and field inoculation were as described in Sermons and Balint-Kurti (Sermons & Balint-Kurti, 2018).

#### 2.1.2 Visual Scoring

Visual scores of disease severity collected at the plot level by expert raters were accepted as the ‘ground truth’ values. Scores from a total of 29 dates were used, averaging roughly 10 per year. All plots were visually scored at least twice and at most five times after anthesis. The SLB scoring rubric is as follows: 9-No evidence of leaf blight; 8-A few spots on the lower leaves; 7-A few spots on the ear leaf; 6-More spots on the ear leaf but the lesions don’t coalesce; 5-Lesions on the ear leaf have grown together, particularly at the tip of the leaf to give quite large necrotic areas; 4-Lesions on the leaf above the ear leaf have grown together too; 3-Leaf above the ear leaf almost completely dead; 2-Almost all tissue on the plant dead; 1-Everything brown (**Figure 1**) (Sermons & Balint-Kurti, 2018). All visual scores were recorded using the Fieldbook app (Rife & Poland, 2014).

**Figure 1:**
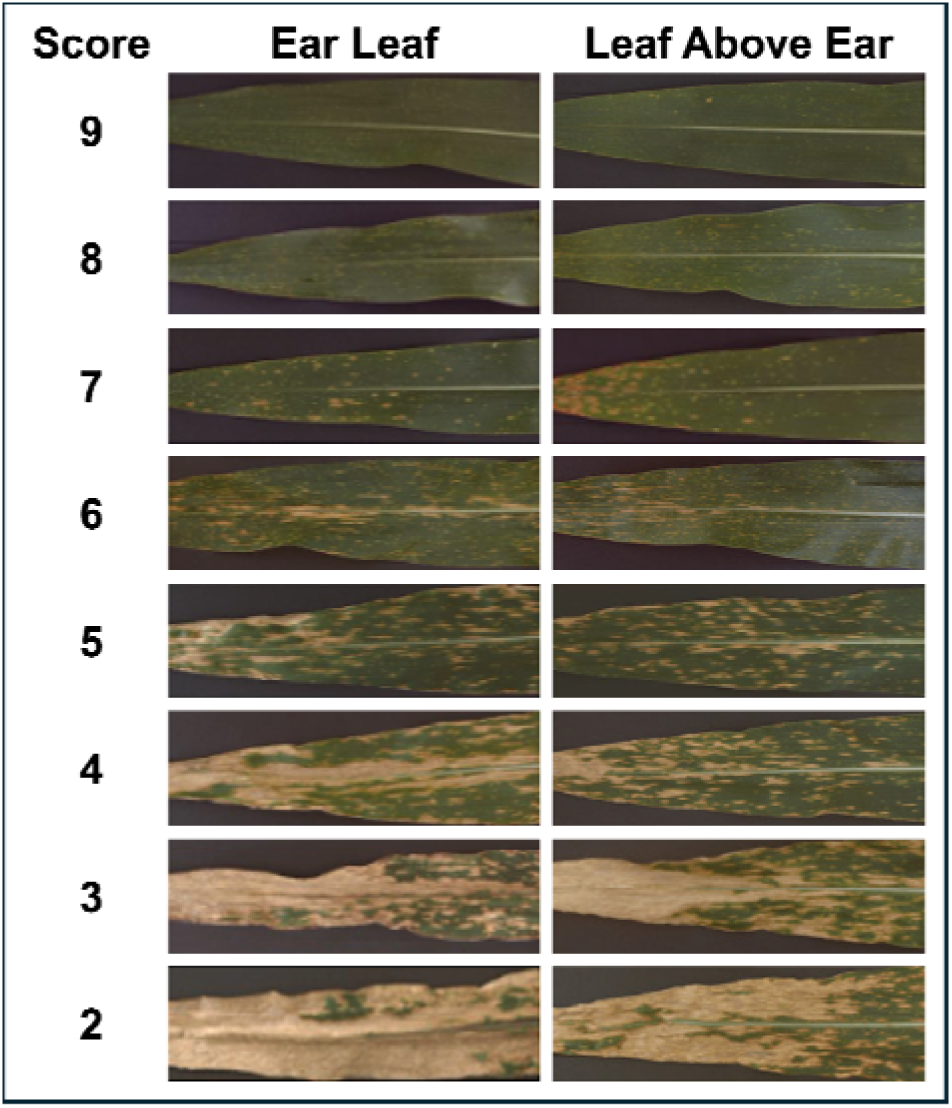
Visual Scoring Guide. The severity score is assigned using a disease diagram to assign scores to each plot with half-integer values used between scores (Sermons & Balint-Kurti, 2018).

### 2.2 RGB SENSOR DATA

#### 2.2.1 Image Data Collection

A DJI Matrice 300 RTK quadcopter and the DJI Zenmuse P1 35 mm full-frame sensor (SZ DJI Technology Co. Ltd.) collected all images. Sensors were selected to maximize resolution, improving the quality of processing and analysis further down the workflow. Flight missions were planned on the DJI Pilot 2 application and were conducted with the following parameters: 80% forward overlap, 80% side overlap, 12 meters (40 feet) of elevation, flight speed of 5.6 kilometers per hour (3.5 miles per hour), and a shutter interval of 1.0 second. These parameters were chosen to balance high ground sampling distance with low flight disturbances (changes in lighting conditions, battery changes). The elevation was the lowest setting allowable in the DJI Pilot 2 application and propeller downdrafts were shown to not significantly affect canopy movement (Tang et al., 2023). Flights were conducted post-anthesis when plants had reached terminal growth.

#### 2.2.2 Image Data Processing

All raw images were constructed into georeferenced orthomosaics using the python API for Agisoft Metashape Pro version 2.1 (Agisoft LLC) using the workflow found at https://github.com/nirwan1265/metashape.git. The coordinate system for the processing and outputs was EPSG:2264 NAD83 / North Carolina (ftUS). A summary of procedure is: (a) RGB images were added to a Metashape project from a folder containing a single flight date; (b) photo alignment was performed with a key point limit of 40,000, tie point limit of 4,000, and downscale of 2; (c) depth maps were built with filter mode set to “Mild” and max neighbors of 100 for depth map generation; (d) point clouds were built with max neighbors of 100 for depth map filtering; (e) classification of ground points with maximum angle of 15.0 degrees, a maximum distance of 1.0 meter (3.28 feet), and cell size of 10 meters squared (107.6 square feet); (f) digital surface models and digital terrain models were built; (g) orthomosaics were built from both model types with blending mode as mosaic, and the refine seamlines option enabled. Digital elevation models (DEMs) and orthomosaics were exported from Metashape as geotiffs.

Plot outlines were drawn as individual polygons for each field using the CreatePlots module from the ArcGIS Pro UAV Toolbox available at https://github.com/reaustin/arcgis-uav-toolbox.git (Austin, 2022/2022). All polygons were labeled with the plot number used during visual scoring using the Sequential Numbering tool in ArcGIS Pro. Each flight’s orthomosaics were then cropped into individual plot images that were named and bound by the shapefile data by using a python script available in GitHub repository at the following link: https://github.com/gagelab/uav_tools.git.

#### 2.2.5 Image Annotation

To use visual scores as the ‘ground truth’ data, UAV images recorded within 3 days before or after a visual scoring event were assigned a label based on the most recent visual scoring. The plot-level .jpg image was given a unique name that identified it based on the necessary information to assign it to a single score: image acquisition date, name of the field site, and the plot number. The images and the information in the unique identifier were stored alongside the most recent visual score within the acceptable day range and the date the score was taken can be accessed at Supplemental Data S1 and https://github.com/gagelab/uav4slb.git (in process; to be finalized before publication).

### 2.3 Model training and evaluation

#### 2.3.1 Model selection

Three families of pre-trained deep learning model architectures were evaluated: vision transforms (ViT), convolutional neural networks (CNN), and hybrids (CNN + ViT). Choices were based on image analysis performance described in the model’s source publication. The three CNNs evaluated were EfficientNet V2-S (Tan & Le, 2021) and two sizes of ConvNeXt V2-B and V2-L (Woo et al., 2023). We evaluated three ViTs: two sizes of DIVOv2 ViT-B/14 and ViT-S/14 (Oquab et al., 2024), and EVA-02-B (Fang et al., 2024a). The hybrid models we evaluated were MaxViT-S (Tu et al., 2022), SwinV2-B (Liu et al., 2022) and CoAtNet2 (Dai et al., 2021). Hyperparameters were manually tuned based on the published findings from each architecture, the nature of our dataset, and available computing capabilities rather than via grid search or other optimization methods.

#### 2.3.2 Model Training and Evaluation

The code associated with model training and evaluation can be accessed at https://github.com/gagelab/uav4slb.git (in process; to be finalized before publication). The total dataset was partitioned into three sets in the following manner: a) a training set with which the model learned image characteristics relevant to the task; b) a validation set for trained model evaluation and adjustment; c) a test set of images that the model has never seen before. We implemented a cross-year evaluation test where data was split into three folds. For each fold, data from two years was randomly split with 85% to the training set and 15% to the validation set. The parameters for the highest performing model from the training and validation phase were saved and then applied to the unseen third year as the test set. We evaluated model performance based on the results from the test set. The loss function for all models was mean squared error (MSE). This is calculated using Equation (1):

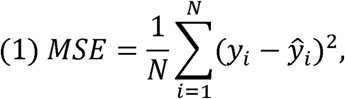

Where *N* is the number of samples, *y_i_* is the visual score, and *ŷ_i_* is the image-based score.

Three metrics were selected to evaluate test performance. The coefficient of determination (R^2^) was calculated according to Equation (2)

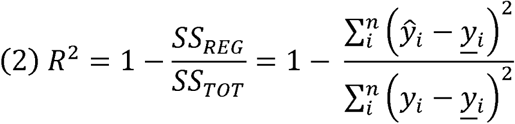

Where *SS_REG_* is the sum of squares regression (variation explained by the model), *SS_TOT_* is the total sum of squares (total variation in the data), *y_i_* is the visual score at observation *i*, y *_i_* is the mean value of all observations, and *y*□*_I_* is the predicted score at observation *i*. The mean absolute error (MAE) was calculated according to Equation (3). The root mean squared error (RMSE) is equal to the square root of the MSE (Equation 1).

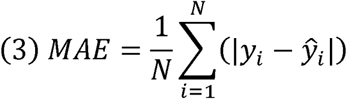

We had the four raters from model training plus one additional rater score SLB severity for 100 plots on the same day, immediately followed by UAV image collection to estimate precision and variability between and among image-derived and in-field visual scores (Supplemental Data S2). From these scores, we calculated the Pearson r correlations for all combinations of rater-to-rater (n = 10) and model-to-rater (n = 5) which essentially treated the best model as a sixth rater.

#### 2.3.3 Sources of error

The circumstances surrounding in-field image collection and visual scoring create diverse sources of error. We categorized sources of error into three levels based on whether they impact an entire flight, a particular image, or an assigned visual score. Flight-level sources of error were conditions affecting spatial and temporal attributes that were maintained throughout the duration of each flight. The image-level sources of error consisted of unique characteristics dependent on location with a field and genotype, making them variable within a given flight and thus measured on a per-image basis. Finally, visual scores themselves were also considered for their contribution to model error due to the tradeoff between dataset size and label noise. The code for organizing and extracting these datasets features can be accessed at https://github.com/gagelab/uav4slb.git (in process; to be finalized before publication).

We explored three types of flight level sources of error: lighting conditions, solar geometry, and crop or disease progression at the time of imaging. Using one-minute radiation observations measured in watts per meter squared (W/m^2^) by the North Carolina Environment and Climate Observing Network station (ECONet) at CCRS (35.66974°N, 78.4926°W), we calculated the irradiance mean and coefficient of variance over the course of all flights (*Cardinal [Data Retrieval Interface]*, n.d.). Using the pvlib python library, we calculated solar zenith angle (SZA), time from solar noon, and shadow-row angle at the midpoint of each flight (Anderson et al., 2023). We normalized the flight dates across the three years for both maize development and disease progress by calculating accumulated growing degree days (AGGD) and days post-inoculation (DPI) per flight. The growing degree units per day were also accessed from the Econet station at the CCRS.

For image-level error quantification, we estimated weed pressure, shadows, brightness, and contrast. To measure per-image weed quantity, we classified all pixels as weeds using a canopy height model to calculate the proportion of each image that was ground (height = 0 feet), maize canopy (height > 2.46 feet [0.75 meters]), and weeds (0 < height ≤ 2.46 feet [0.75 meters]). For per-image shadow fraction, image pixels were converted from the RGB (Red - green - blue) to HSV (Hue - saturation - value) color space and classified using a qualitatively derived V-channel (brightness) threshold, normalized to the 95% percentile of whole-image value. Mean image brightness was calculated for each image by averaging the V-channel for all pixels in the HSV color space to estimate the effect of overall light intensity on a per image basis. Contrast root mean square (RMS) was calculated as the standard deviation of pixel intensity after converting images to grayscale to capture the variation of perceptually weighted luminance.

Association of visual scoring noise and error was estimated with the number of days between UAV flights and visual scoring (flight-score offset). We calculated the signed error as the test score minus the visual score then correlated the difference across the seven day-wide windows used for image labeling. Additionally, we discovered plot images that show a departure of viewable disease characteristics from the corresponding visual score used during training and testing.

## 3 RESULTS & DISCUSSION

### 3.1 Dataset Characterization

#### 3.1.1 Image Count Distributions

The visual scores remaining after filtering corresponded to 26,071 plot images and covered the full one-to-nine range of SLB severity (**Figure 2A**). Interestingly, the proportion of whole integer scores used was 63.6% while only 36.4% of the scores assigned were half-integer values (**Figure 2A).** We fit a kernel density to the visual scores to provide a corrected, non-uniform expected count for each of the 17 scores that accurately reflects the underlying distribution of observed severity. Integration of the kernel density estimate with ±0.25 windows lead to a ∼50% proportion of half to whole integer scores for any smooth distribution over the one-to-nine SLB scoring scale. The observed proportion of half-integer scores fell well below the expected proportion and a chi-squared goodness-of-fit test confirmed systematic departure across all 17 values (χ²(16) = 1,915.44, p < 0.001). This is consistent with the tendency for raters to assign the scores pictured in the rating protocol, a behavior known as “knotting” (Bock et al., 2008; Schwanck & Del Ponte, 2014; Sherwood, 1983). This phenomenon can be considered a tradeoff between ease of use and score resolution. The familiarity granted by a visual guide such as **Figure 1** increases the convenience and perceived speed of rating (Madden et al., 2017). On the other hand, the challenge of interpolation limits scorers from assigning intermediate values (Large, 1966). These findings motivate image-based automation as a route to regain resolution in disease severity assessment.

**Figure 2:**
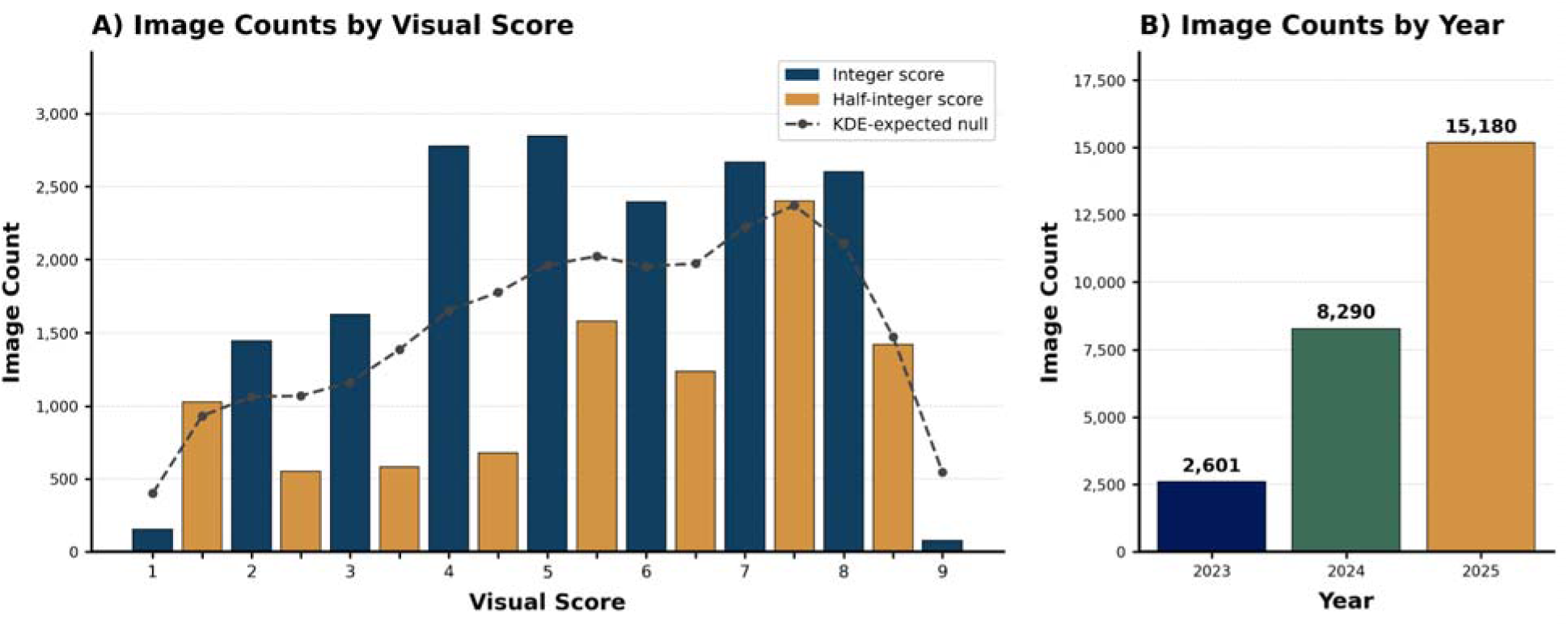
Distribution of visual scores and quantity of scored plot images. (A) Counts of images by corresponding Southern leaf blight (SLB) visual severity score with 9 indicating a plot with no evidence of leaf blight and 1 indicating everything brown. Half-integer scores in orange and integer scores in blue. The dashed line represents the expected score frequency derived from a kernel density fit to the observed data, reflecting the non-uniform underlying severity distribution. (B) Counts of filtered image-visual score pairs per year with 2,601 images in 2023 (blue), 8,290 images in 2024 (green), and 15,180 images in 2025 (yellow).

The number of images with corresponding visual scores varied considerably from year to year, with 10% recorded in 2023, 31.8% from 2024, and 58.2% from 2025 (**Figure 2B).** This imbalance was a result of improvements of protocols over the course of the experiment, such as beginning flights earlier in the season and increasing the frequency of flights (**Figure S1**). A combination of these factors may contribute to the significantly different distribution of disease scores between years (**Figure S2).** Summary statistics of visual scores for each year are included in **Table S1.**

### 3.2 Model Performance Evaluation

#### 3.2.1 Year-To-Year Generalizability

One of this study’s objectives was to evaluate a variety of deep learning models for applicability in prediction problems pertinent to maize genetics and breeding programs. Nine models were selected from three architecture families based on ability to predict from fine-scale image features. Our model selection process was rooted in the biology of the trait of interest, aiming to stimulate the discernment of disease characteristics without explicit annotation. The systematic differences between each year’s imaged experiments influence both this biology and perception of disease severity through variation in conditions such as soil type, weather conditions, germplasm, and weed pressure (Ullstrup, 1972). The actual feasibility of an image-based pipeline to alleviate the labor requirements of SLB scoring necessitates performance evaluation on field experiments that were not used during model training.

We implemented a 3-fold cross-year evaluation test by allocating two years of plot images and visual scores for model training and one year for model testing (**Table 1**). Such evaluation simulates a major barrier to the deployment of image-based agronomic models: certainty that test scores are derived from features relevant to the target rather than those distinct to a particular environment (Jarquín et al., 2017). For each fold, models were trained on two years then used to estimate disease severity in plot images from the target left-out year to test year-to-year generalizability. By isolating the left-out year’s unique growing and scoring conditions from the training data, we aimed to reduce inflation of model test performance due to shared conditions in both the training and testing data. Additionally, this approach emulates domain generalization, a technique that improves performance reliability on test data by training on multiple source distributions (Hu et al., 2025). Models with high test performance are assumed to have learned the image features relevant to disease severity despite the different growing environments presented during training versus testing.

**Table 1:**
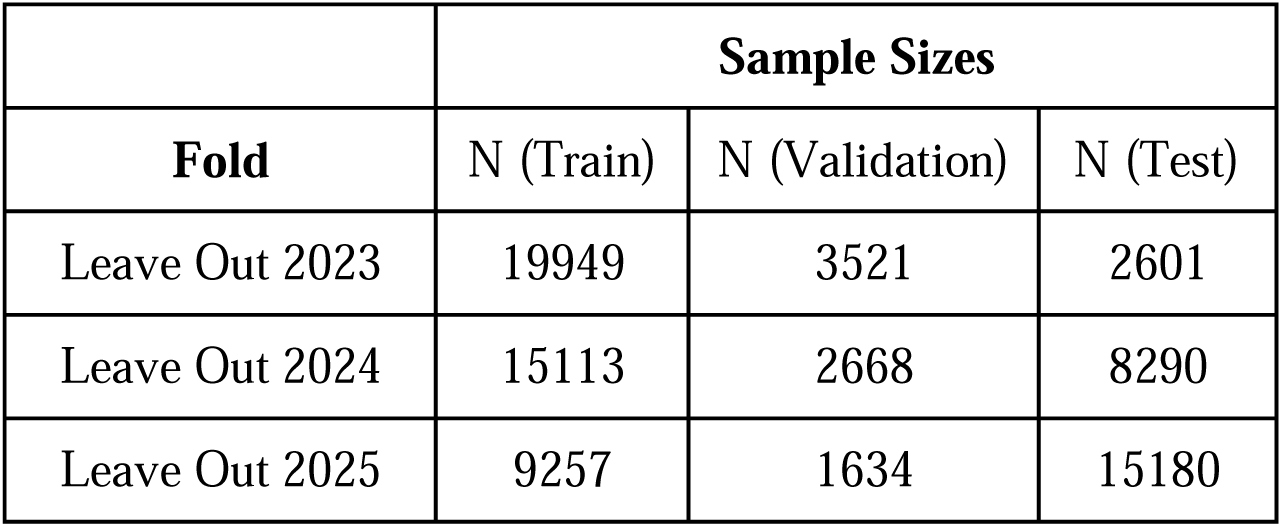
Number of images per split in cross-year test evaluation folds. One year was excluded from training and reserved for the test set. The images from the remaining two years were pooled together and randomly split 85% into training and 15% into validation.

|  | Sample Sizes |  |  |
| --- | --- | --- | --- |
| Fold | N (Train) | N (Validation) | N (Test) |
| Leave Out 2023 | 19949 | 3521 | 2601 |
| Leave Out 2024 | 15113 | 2668 | 8290 |
| Leave Out 2025 | 9257 | 1634 | 15180 |

We tested nine different deep learning models, each of which had distinct combinations of architecture and pre-training datasets. We selected R^2^, MAE, and RMSE as performance metrics and averaged these across all three folds during model comparison. Across all metrics, EVA-02-B, a pre-trained vision transformer with a DINOv2 backbone, achieved the highest performance (Fang et al., 2024b) (**Figure 3A-C)**. While EVA-02-B achieved only the third highest R^2^ when 2023 was withheld as a test set, improved performance on the 2024 and 2025 test sets compensated for this deficiency (**Figure S6**). One possible contribution to lower performance on the 2023 test set was the unbalanced nature of the dataset as described above. The number of images in the 2023 test set (2,601) was 5,689 fewer images than 2024 and 12,579 fewer than 2025. With less image feature variation captured in a smaller test set, the complexity of spatial relationships learned from a highly variable training set may have limited model performance (Hu et al., 2025).

**Figure 3:**
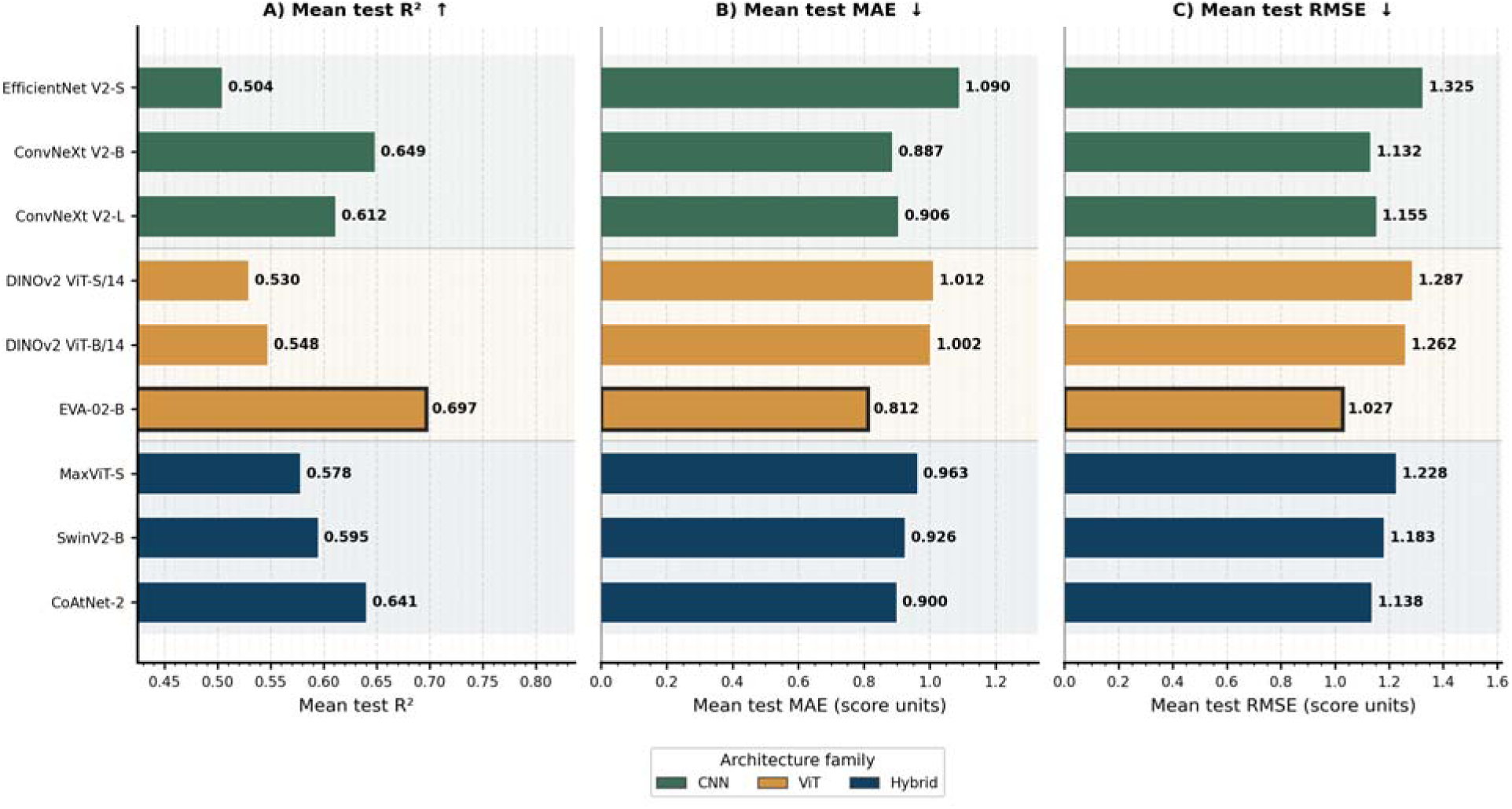
Average model performance metrics in cross-year test evaluation organized by architecture family. Bold outline indicates best performance. A) Average of coefficient of determination R^2^ values from all test sets; high values indicate better performance, symbolized with an up-arrow. B) Average mean absolute error (MAE) from all test sets; low values indicate better performance, symbolized with a down-arrow. C) Average root mean square error (RMSE) from all test sets; low values indicate better performance, symbolized with a down-arrow.

After three iterations of training and testing, the model-estimated scores for the test years from all folds were compiled and fitted to a regression line using ordinary least squares (OLS) to determine the relationship between the test scores and visual scores. The slope of the regression line was 0.76, demonstrating a slight compression towards the middle of the scale (**Figure 4A**). This was further supported by the increase in mean signed error at the extremes of SLB severity, which shows systematic regression of test scores towards the training mean and away from the distribution tails (**Figure 4B**). One potential explanation for this is the lack of available training data due to low frequency of visual scores at the ends of the scale. Additionally, the high-resolution details capture disease severity in the combination of healthy and necrotic tissue from images with mid-range scores. Model-estimated scores at the extremes may be less reliable due to the lack of separable necrotic and healthy tissue: a high visual score image typically contains only full, green and healthy plants while those with low visual scores contain a vacant, brown, and dead plot.

**Figure 4:**
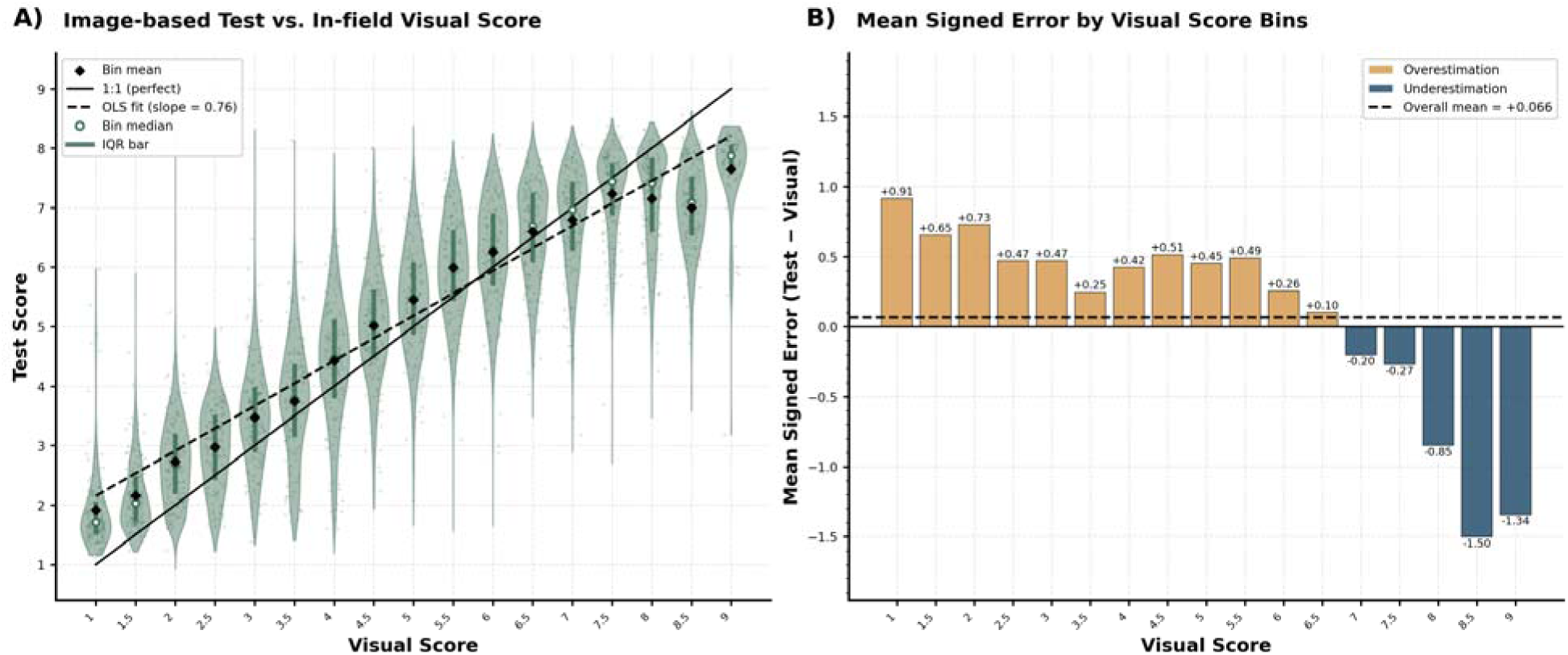
Test performance of combined folds. A) Violin plots by 17-bin visual scoring scale. Green bars represent the inter-quartile range. Black diamonds represent the bin mean and white dots represent the bin median. The solid black line is the 1:1 reference line and the dashed black line is the ordinary least squares fit line (slope = +0.76) of all imaged-based test scores to rater-derived visual scores. B) Bar chart of mean signed error by 17-bin visual scoring scale. The yellow bars represent an overestimation of disease resistance (positive error) and blue represents an underestimation of disease resistance (negative error). The dashed black line represents the overall mean signed error (+0.066).

#### 3.2.2 Comparison of rater-to-rater and model-to-rater correlations

Variability between expert human raters scoring the same plots under identical conditions can lead to discrepancies in the identification of small effect QTL (Poland & Nelson, 2011). In our SLB visual scoring protocol, one rater is encouraged to collect all ratings for a particular study which improves consistency within an experiment. Where visual scoring may be lacking is reproducibility across multiple experiments. The plot-level images and scores used in training and testing were taken from multiple experiments, resulting in visual scores from four different raters (listed in the Acknowledgements section). In this way, we aimed to develop a model that can address this drawback of visual scoring by uniformly estimating severity across multiple experiments with an automated approach.

To test this, we had the four raters from model training plus one additional rater score SLB severity for 100 plots on the same day, immediately followed by image collection to generate image-based test scores that we used to evaluate whether the model agrees with raters as well as raters agree with each other. A pairwise Pearson r was calculated for all rater-rater combinations (n = 10) to estimate rater-to-rater variability. We then calculated pairwise Pearson r for all model-rater combinations (n = 5) to assess model-to-rater variability, essentially treating the model as a sixth rater. The mean of rater-to-rater correlations was 0.80, ranging from 0.69 to0.86 (**Figure 4**). The mean of the model-to-rater correlations was 0.75, ranging from 0.70 to 0.80. The model-to-rater correlations were on average lower than the rater-to-rater correlations by 0.05. The greater range in rater-to-rater correlations (0.17) than model-to-rater correlations (0.10) indicated greater variability from visual scoring. Raters who assigned similar versus dissimilar scores have the correlations at the higher and lower ends of the range. This was driven by rater bias in the visual scores, indicating that model derived scores may offer comparable precision. These results substantiate the utility of automated phenotyping using models trained from multiple raters without need for repeated scoring. The process of annotating large drone image datasets with accurate labels is a major barrier for model development (Guo et al., 2024; Wang et al., 2026). The approach used in this study suggest that individual labels derived from different experts can be pooled during training to efficiently create models that address both subjectivity and labor intensity of in-field disease assessment and image dataset curation.

### 3.3 Sources of model error

#### 3.3.1 Flight conditions explain model error

Variations in lighting conditions, solar geometry, and time of year represent powerful domain shifts influencing deep learning prediction outcomes in agriculture imagery (Chakhvashvili et al., 2024; Hu et al., 2025; Spisak, 2022; Valencia-Ortiz et al., 2021; Zhu et al., 2024). These features introduce inconsistencies that are intrinsic components of passive remote sensing, adding complexity to machine learning tasks. We evaluated the robustness of EVA-02-B to known and hypothesized sources of error from flight-level conditions by computing Spearman correlations (11) to characterize associations to EVA-02-B per flight MAE; = 0.05 was used as a significance threshold given the small sample size (n = 27 flights) and exploratory nature of the analysis (Figure 5). Values mentioned in this section are shown with higher precision in Table S2, along with the mean and standard deviation of each flight characteristic. This analysis dissected the strengths and limitations of our approach, providing insights on both the applicability of current procedures as well as targets for potential improvement.

**Figure 5:**
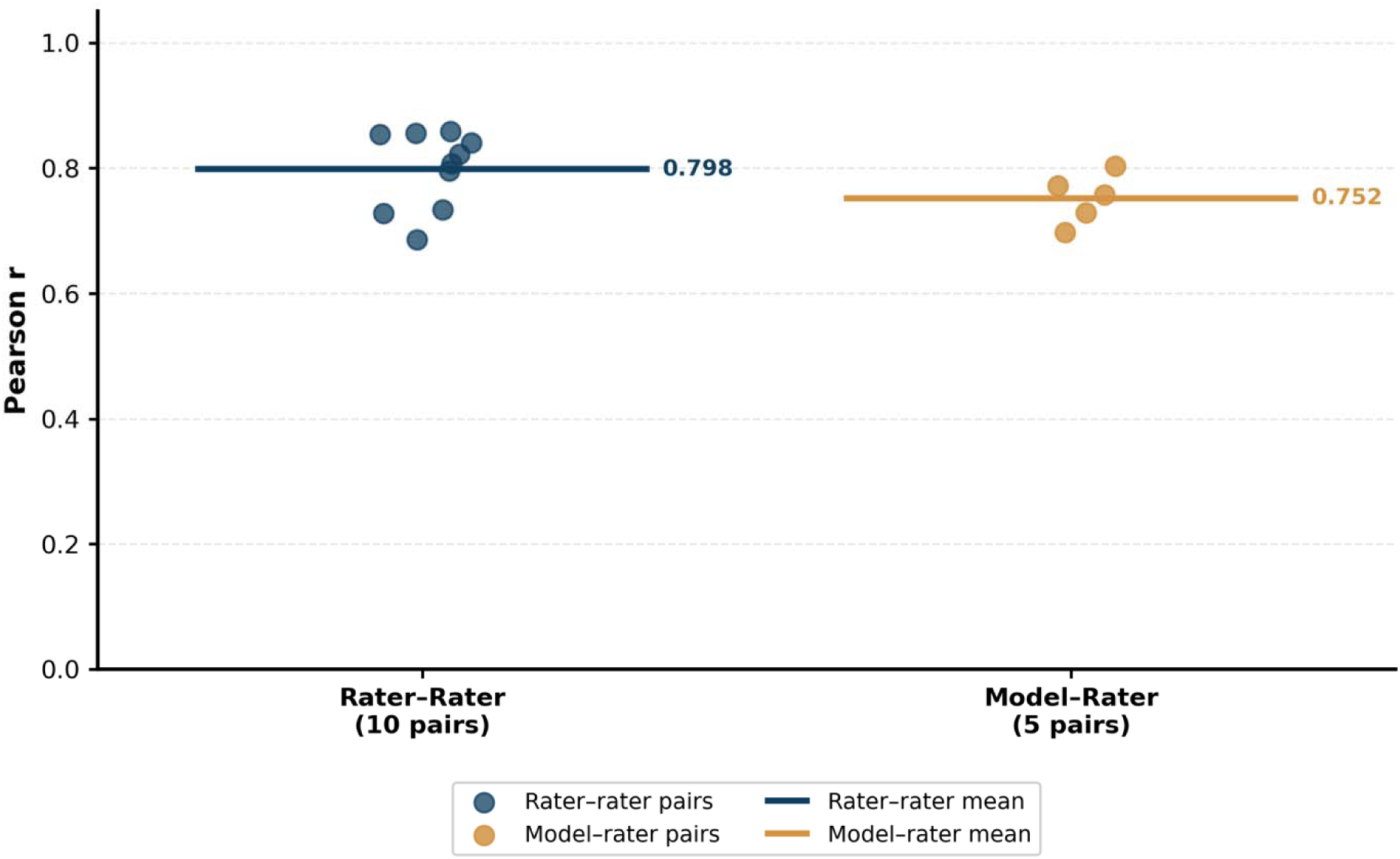
Scatter plots of rater-rater Pearson r (blue) and model-rater Pearson r (orange). From five raters, the 10 rater-rater Pearson r values had a mean of 0.798 and the 5 model-rater Pearson r values had a mean of 0.752. The mean values are represented in the figure by horizontal lines. The difference between these means was 0.046.

Flight-level lighting conditions are a core consideration for agriculture remote sensing as light intensity and variance have been shown to influence highly informative spectral and textural features of collected images (Bongomin et al., 2024; Chopin et al., 2018). We calculated the irradiance mean and coefficient of variation (CV) over the course of all flights (North Carolina State Climate Office, n.d.). The association between per-flight mean irradiance and MAE was statistically significant (p = 0.015) and had a negative effect size (11 = −0.463). The irradiance CV had a weak positive correlation (11 = 0.205) to MAE that was not statistically significant (p=0.305).

Another category of interest was the solar geometry conditions for each flight. Solar zenith angle (SZA) is the angle between incoming radiation and a line perpendicular to the plane of the field; as SZA increases, so does shadow length (Li et al., 2025; Valencia-Ortiz et al., 2021). Time from solar noon is the absolute difference in minutes between flight start time and the daily solar noon (the moment at which the sun reaches its maximum elevation angle at a given location) which varies by geographic position and day of year. Best practices for UAV imaging recommend flying at solar noon to minimize the bidirectional reflectance distribution function and the length of shadows (Bongomin et al., 2024; Shahi et al., 2022). The taller canopy of maize caused shadows to cast within a plot or onto neighboring plots, leading to inconsistent illumination. The extent of this problem depended on the SZA and the angle of the incoming radiation relative to plot orientation. We calculated shadow-row angle for each flight to capture shadow direction, ranging from perpendicular (90°) to parallel (0°) with the row. We found no statistically significant association between any of the three measurements of solar geometry and EVA-02-B MAE.

To assess the effect of in-season flight time, we normalized the flight dates across the three years for both maize development and disease progress by calculating accumulated growing degree days (AGGD) and days post-inoculation (DPI) per flight, respectively. We found AGGD (p = 0.001, IZ = 0.590) and DPI (p = 0.009, IZ = 0.490) had a statistically significant and large effect size on EVA-02-B MAE. These results suggest that errors track crop phenology and disease symptomology more than pure image quality variation from lighting and solar geometry. As the plants and disease progress through the season, two different sources of necrotic tissue emerge: SLB infection and senescence. Failures in the model to distinguish the two sources of necrosis may be leading to greater errors later in the season.

#### 3.3.2 Model robustness to image noise

In remote sensing data, background complexity, illumination variation, and shadow effects contribute noise to each image. We randomly chose the high-error examples from the fourth error quantile (greater than the 75th percentile) for qualitative analysis (Figure 7). We asserted that weeds, shadows, contrast, and brightness may be sources of noise contributing to errors. While there are instances when EVA-02-B made accurate predictions despite high levels of noise, there were consistencies amongst those images with high prediction error that we used to justify further analysis. The amount of weed pixels varied spatially and temporally by timepoint and field while shadows, contrast, and brightness varied within each flight (**Figure S4**). We aimed to evaluate measurements of these sources of image noise to determine their relationship to per-image error. This analysis could be used to inform strategies to strengthen model performance and generalizability such as field management procedures, flight planning targets, and data stratification methods.

All image features tested had a statistically significant though weak relationship with absolute error. Weed fraction and contrast RMS had the highest albeit weak correlations with absolute error at 0.119 and −0.123, respectively (**Figure 8A-D**). This implies that a diverse range of image characteristics were available during training that strengthened model year-to-year generalizability. Brightness and shadows are known sources of noise in remote sensing in agricultural research yet our deep learning approach demonstrated resilience to these (Mardanisamani & Eramian, 2022; Shahi et al., 2023). While weed fraction and contrast RMS are weak drivers of model error, both have potential to be addressed in preprocessing steps.

#### 3.3.4 Label noise

High-quality image labels are critical for training effective models. Expert raters score SLB severity longitudinally to calculate the area under the disease progress curves (AUDPC). This AUDPC value serves as the phenotype for most genetic studies focused on maize disease resistance. In our study, we required no additional time commitment from expert raters to annotate lesions, score images, or any other form of training data labeling. Instead, we appropriated these visual scores for image labels, greatly streamlining the process of generating training data. A drawback of this application of in-field visual scores is the discrepancy in frequency between scoring and imaging (**Figure S1**). We accepted an offset between scoring and imaging of up to three days as a necessary tradeoff between dataset size and label quality. For instance, the majority of images acquired in 2024 received labels offset by three days. If we reduced the maximum offset to two days rather than three, the 2024 image count would have dropped by 59% and the overall image count by 20% (**Figure S5**). Disease severity is a phenotype that can progress temporally, and we acknowledge that the three-day span is inexact. However, we considered the detrimental effects of the year imbalance and shrunken dataset to be greater than the label noise introduced by a three-day flight-score offset.

With this accepted range, plot images generated from one flight may receive visual scores recorded across a window as wide as up to seven days. We hypothesized that this flight-score offset introduced a systematic bias in signed error as images acquired before visual scoring capture plants with less disease relative to those plants observed days later by expert raters, indicating that EVA-02-B perceived disease progression over a maximum of three days in this dataset. The relationship between signed error and flight-score offset was found to be statistically significant with a weak negative correlation (p < 0.001, 11 = −0.1648). As expected, test scores from images acquired before visual scoring tended to underestimate disease severity and those from images acquired after scoring tended to overestimate disease severity (**Figure 8**). Over the course of the imaging period, perceived severity increased due to both disease progression (more necrotic lesions) and maize phenology (more senescence). Aerial images captured prior to scoring showed plots with less disease and senescence than was manually observed in the same plot one to three days later. On the other hand, images acquired after scoring contained more disease-relevant details relative to what the expert raters had observed in the days prior. Thus, the test scores produced from RGB aerial imagery by EVA-02-B seemingly accounted for fine-scale changes in disease severity that occurred during the days between image capture and visual scoring.

The impacts of plant architecture on the perception of disease lesions within the canopy are understandably not equivalent between the image-based and visual scoring methods (**Figure 10)**. Disease assessments taken in-field have an advantage over processed aerial imagery in that the full canopy is viewable from multiple angles. On the other hand, orthomosaics provide only the most overheard nadir perspective where leaf dimensions and orientations affect visibility into\ the canopy. This gives expert raters significantly more visual signals of disease severity to incorporate into their decision when scoring plots. When evaluating scenarios of high absolute error, we found instances where the model-derived predictions appeared to more accurately reflect the disease symptoms depicted in the image than the visual score (Figure 10). This suggests a key limitation of visual scores as labels for aerial images as the manifestation of quantitative disease resistance phenotypes extends beyond the dimensions of the input features.

Drone image processing and orthomosaicing itself generates artifacts under standard field conditions due to canopy movement and plot arrangement. This further complicates the separation of plot mislabeling from true model error as raw image features otherwise cohesive with the visual score are distorted after processing. Label noise due to discrepancies between image and visual score data represents the critical tradeoff between accuracy and throughput. As we attempt to phenotype a complex trait using massive amounts of incomplete image data, the loss of information and its relationship to acceptable performance must be contextualized in a task-specific manner. Furthermore, our ability to assess this tradeoff is partially confounded by the assumption that the visual scores are a perfectly accurate ground truth (Figure 10).

### 3.4 Conclusion

This study investigated the capability of deep learning models to automate the estimation of maize quantitative disease severity from UAV imagery. Over three growing seasons, we developed a dataset of 26,071 plot images labeled with in-field visual scores taken by expert raters. We evaluated nine deep learning models with an agronomically relevant cross-year evaluation to test generalizability onto future growing seasons. By treating visual scores taken on a one-to-nine scale in increments of 0.5 as quantitative and continuous values, we used standard regression model evaluation metrics to determine the highest performing model. We compared correlations amongst expert raters to correlations with image-derived scores and ascertained the model effectively estimated disease resistance with the advantage of comparable accuracy and substantially greater throughput.

The model errors were analyzed against potential contributing factors from UAV flights, individual plot images, and the visual score labels. The mean absolute error per flight showed weak correlations to solar geometry, lighting conditions, and in-season timing (Figure 6).

**Figure 6:**
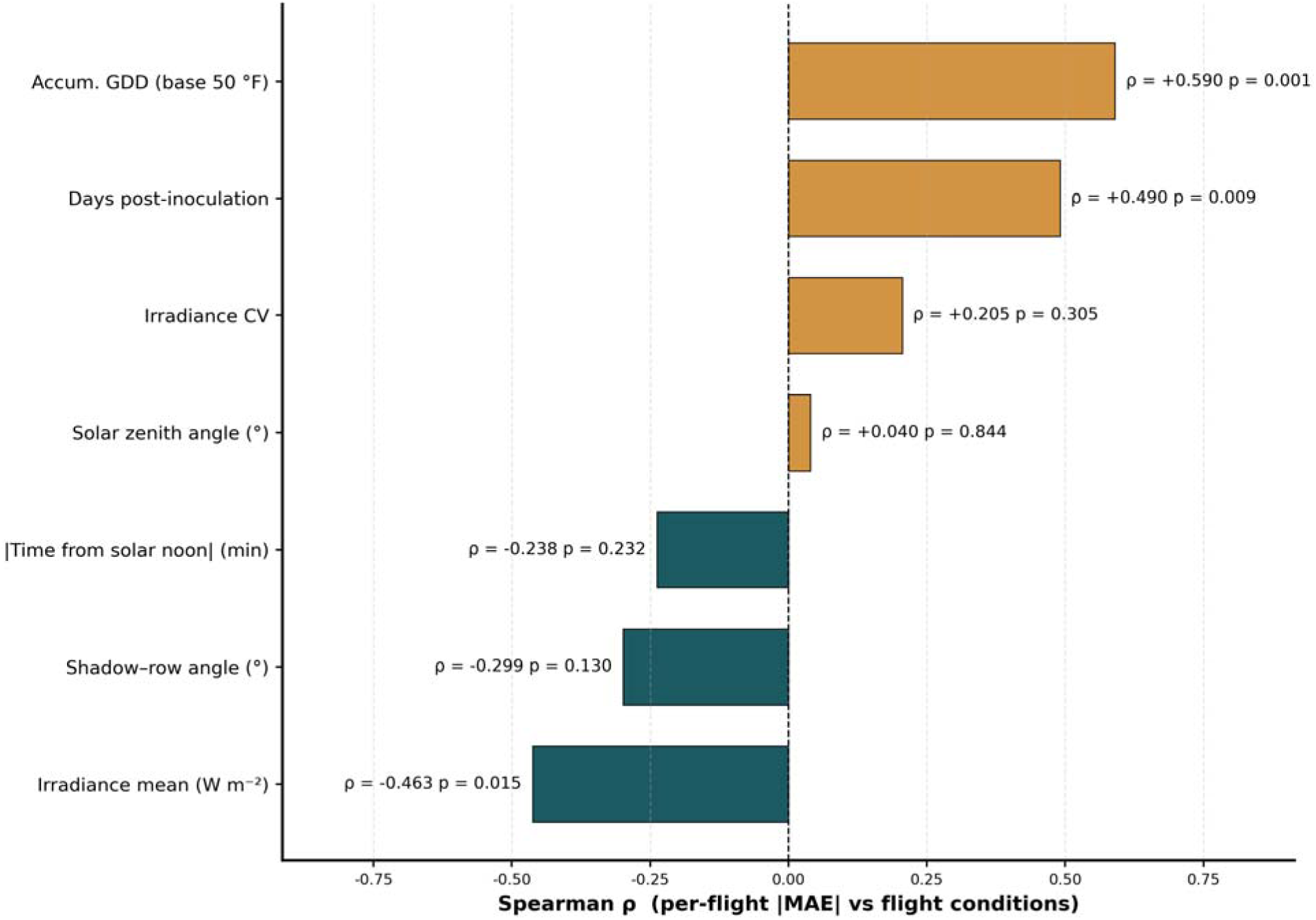
Spearman correlation between flight-level conditions and EVA-02-B mean absolute error. Flight-level conditions evaluated fell into categories of lighting (irradiance mean, irradiance coefficient of variation), solar geometry (time from solar noon, shadow-row angle, and solar zenith angle), and temporality (accumulated growing degree days [AGGD], days post-inoculation [DPI], and flight-score offset). Mean irradiance (p = 0.015), AGGD (p = 0.001), and DPI (p = 0.009) were found to be statistically significant. Mean irradiance had a medium effect size (IZ = −0.463) while AGGD (IZ = 0.590) and DPI (IZ = 0.587) had a large effect size.

**Figure 7:**
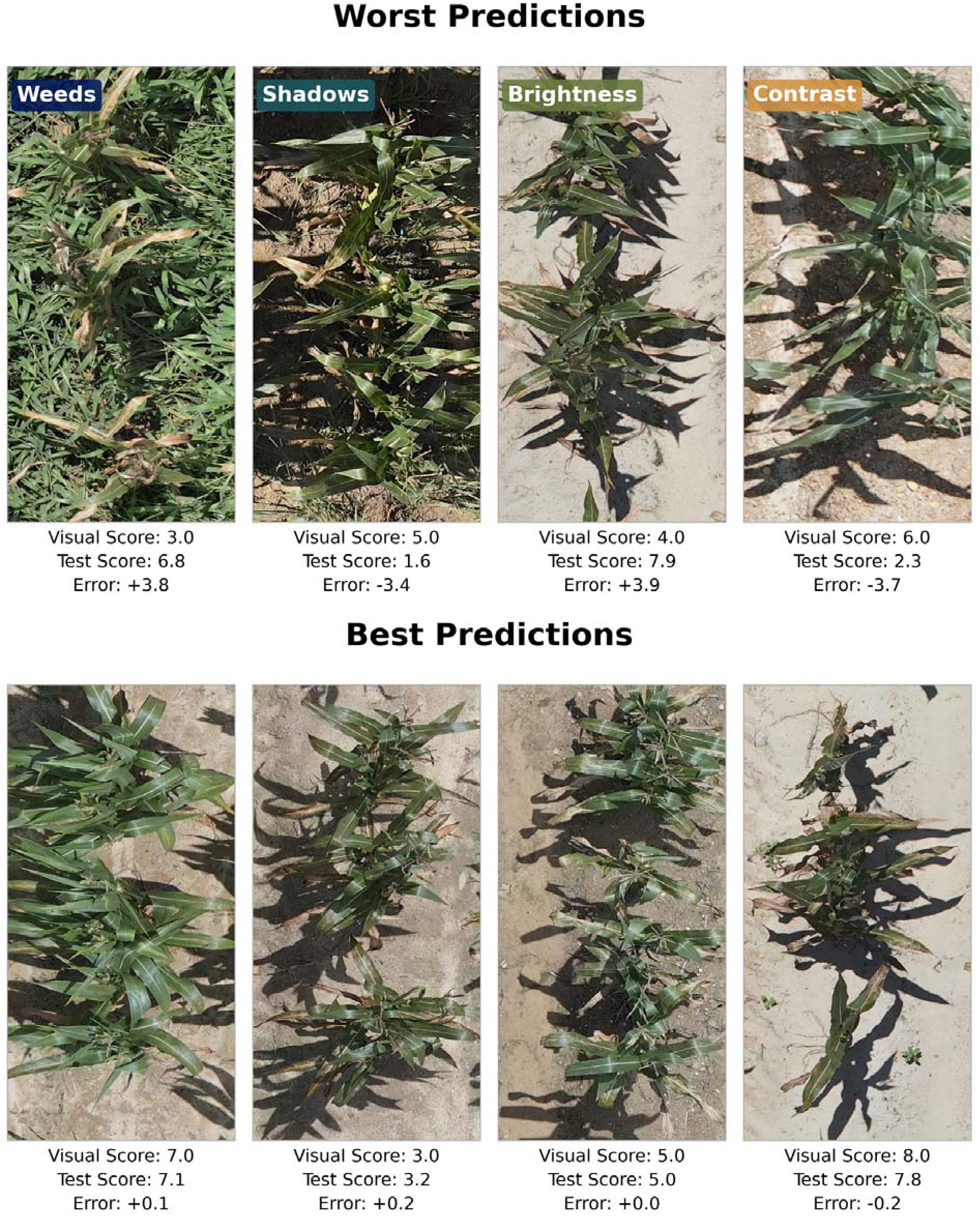
Example images with high (top row) and low (bottom row) absolute error. High error plots are labeled with qualitatively observed image features that may potentially contribute to error.

Absolute error also proved to be robust to image-level features with weak correlations to proportion of weeds, proportion of shadow, mean brightness, and contrast (Figure 8). Label noise introduced by the date misalignment between visual scoring and imaging showed an error pattern indicating our approach achieved fine-scale monitoring of disease progression over time (Figure 9). Thus, we conclude that changes in perceived resistance during the days of separation between visual scoring and aerial imaging is a key source of model error from label noise. Our approach to image dataset creation successfully captured the high diversity of field conditions present in agronomic research. When analyzed with pre-trained deep learning models, the outcome was high-quality predictions, shown to be based on disease-relevant details and proven to be robust to various sources of noise.

**Figure 8:**
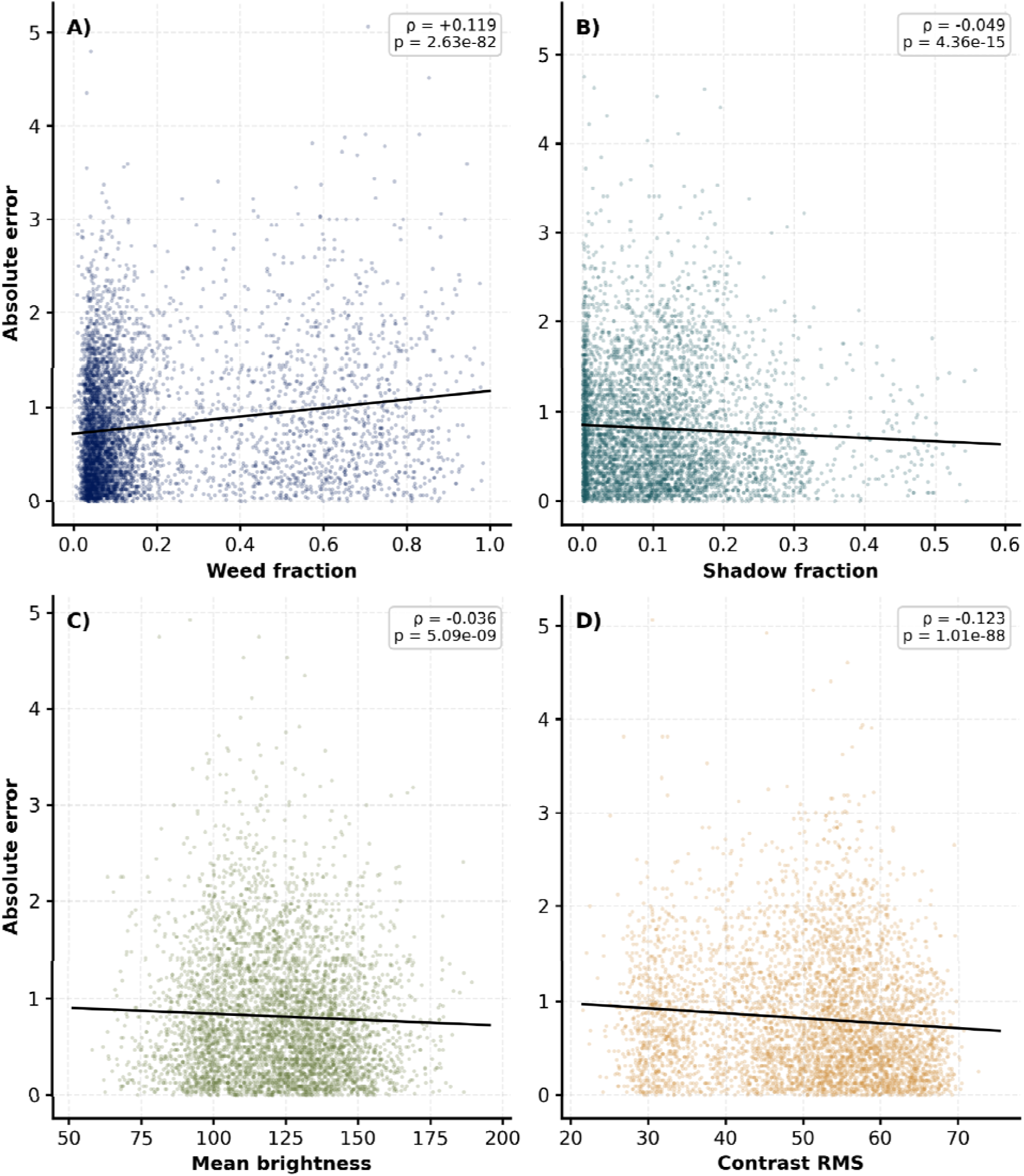
Scatter plots of absolute error and image-level characteristics. Fitted ordinary least squares regression represented by the solid black line. A) Fraction of pixels classified as weeds based on height. B) Fraction of pixels classified as shadow by thresholding pixel values for brightness normalized per image after conversion from red–green–blue (RGB) to hue–saturation–value (HSV) color space. C) Mean brightness as the average of all pixel values after conversion from RGB to HSV color space. D) Contrast root mean square as the standard deviation of pixel intensity after converting images to grayscale to capture the variation of perceptually-weighted luminance.

**Figure 9:**
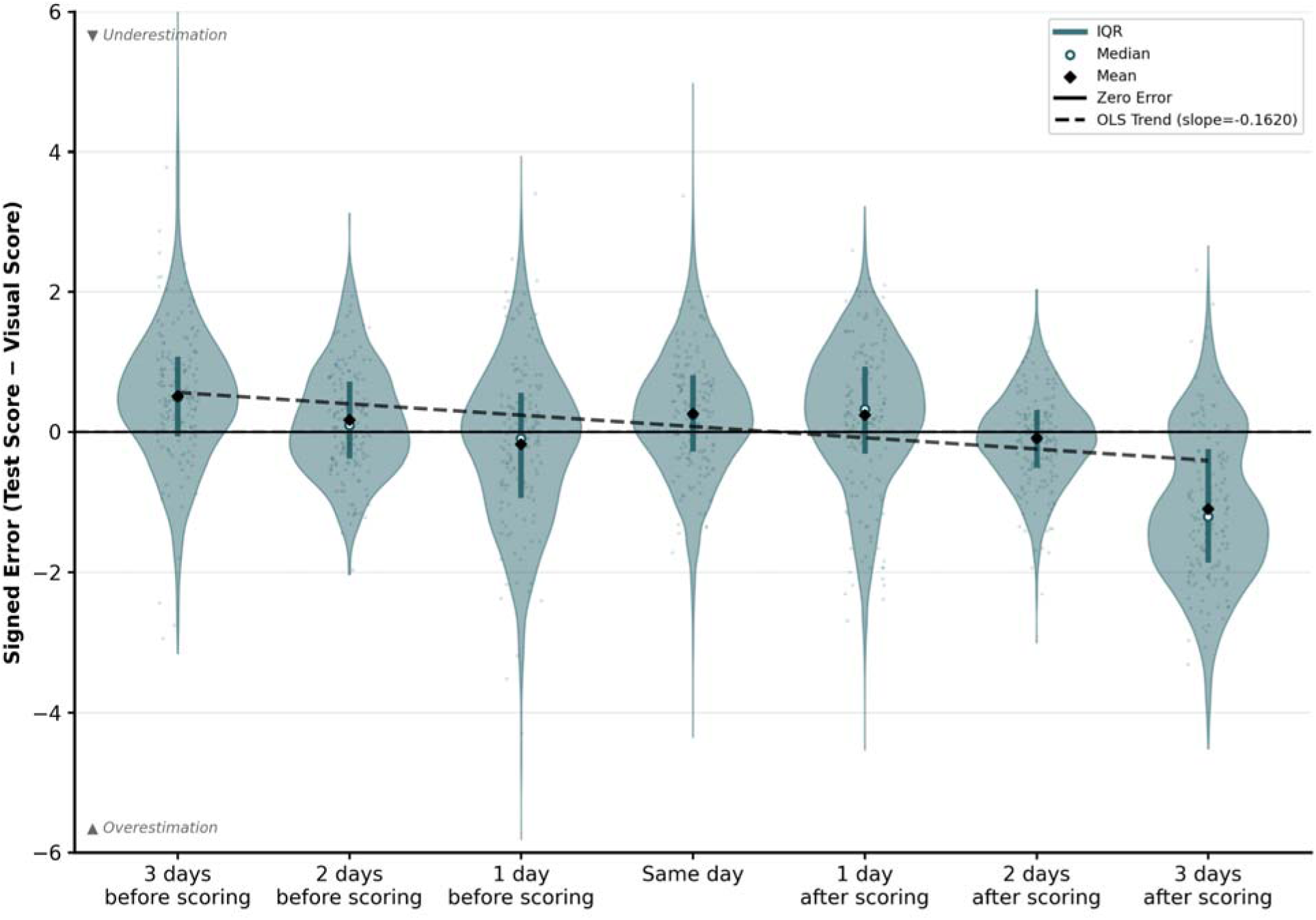
Distribution of per-plot signed error for the number of days difference between visual scoring and imaging. Median and mean are shown as a white dot with teal outline and a black diamond within each violin, respectively. Inter-quartile range (IQR) is shown as the blue line within each plot. Zero error is represented by the gray dashed line. Ordinary least squares (OLS) trend line is depicted in dark red (slope = −0.1620).

**Figure 10:**
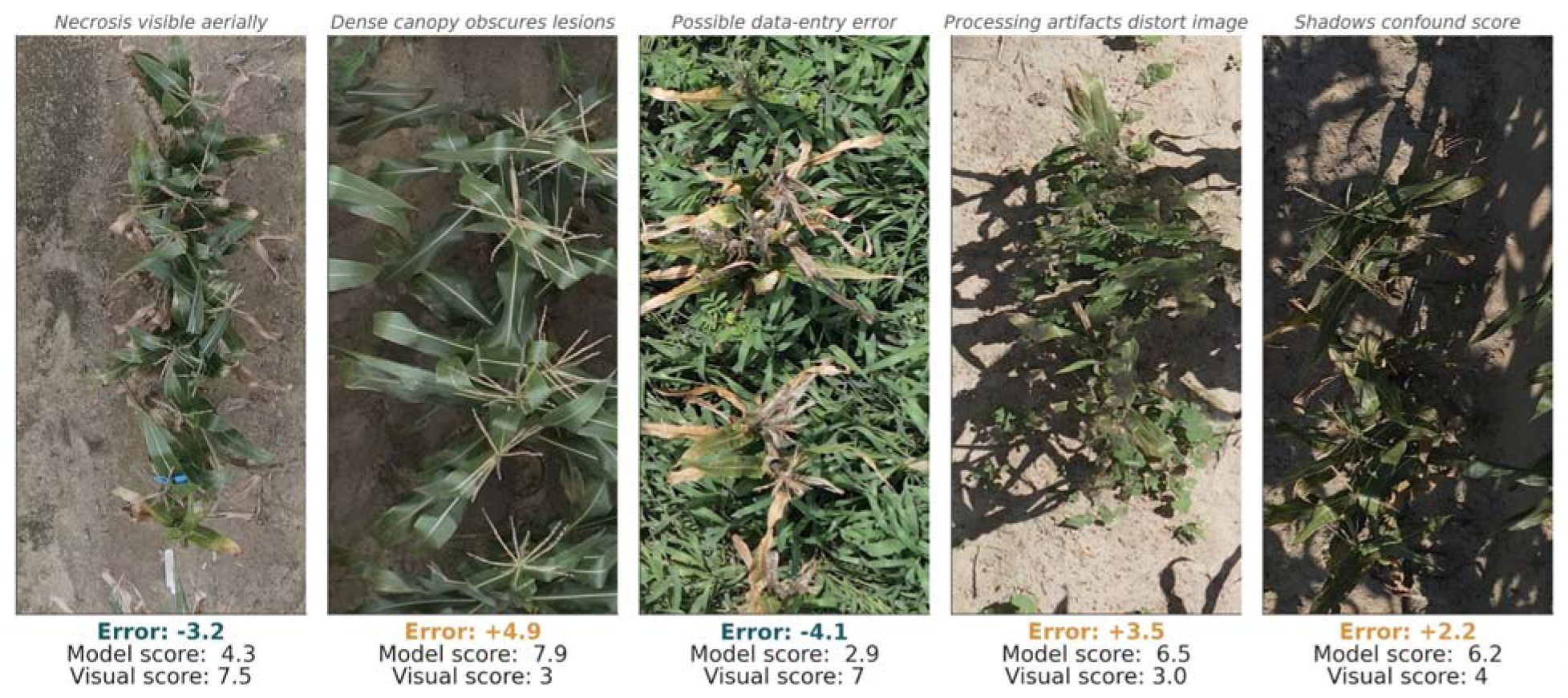
Examples of mismatches between plot image and visual score with proposed specific driver of error.

Phenotypic attributes beyond disease resistance can be extracted from the high-resolution RGB image dataset developed for this study. Leveraging the wealth of information available in these images allows additional analysis of phenotypes with known relevance to disease severity, such as leaf angle and plant maturity. Advancements in deep learning algorithms for image analysis empower the discovery of novel variables from our image dataset. Unsupervised learning, a familiar statistical method for revealing hidden patterns in data, can be implemented in the context of our study to discern previously unknown relationships in disease resistance phenotypes. Self-supervision for deep learning image analysis learns rich representations that can both resolve latent phenotypes from image data and provide pretraining for downstream supervised learning.

This study found promising results that lay groundwork for improved phenotyping of maize quantitative disease resistance. We assert that our approach can alleviate the barrier to experiment size and consistency imposed by in-field visual scoring for SLB resistance. Since each flight captured images of thousands of plots in less than one hour, disease severity can be estimated over larger fields and with higher frequency than is possible with visual scoring. This increase in scoring frequency paves the way for future studies on the genetic mechanisms underlying variation in the temporal dynamics of SLB disease resistance (Alladassi et al., 2026). The increase in scorable field size has potential to strengthen the statistical power of genetic studies when larger populations and more repetitions can be included. The actual feasibility of our approach could be studied with downstream validation benchmarks such as disease resistance QTL identification with model-derived scores. Such future work would solidify HTP as an effective method for genetic dissection and integration of QDR into elite maize hybrids.

Our extraction of underlying mechanistic variables in image processing and analysis provides a procedural basis for improved modeling via multimodal data fusion, informed model-error correction, and guided ensembling (Ispizua Yamati et al., 2026).

## ACKNOWLEDGEMENTS

Visual scores used as image labels were provided by Asher Hudson, Shivreet Kaur, and Tao Zhong. Plot-level image cropping from orthomosaics was done with modified code originally written for the Gage lab by Lucas Bauer. Orthomosaicing pipeline was done with code provided by Nirwan Tandukar. We thank Greg Marshall, Shannon Sermon and the staff at the Central Crops Research Station for assistance with field trial design and management. Loaner equipment provided by Robert Austin.

## SUPPLEMENTAL MATERIAL

Figures in this section are as follows: S1) a visualization of image counts by score and flight dates; S2) violin plots of scores by year; S3) horizontal bar chart R^2^ values for each model by test year; S4) two images of orthomosaics demonstrating data noise potentially driving error; S5) the image counts per year by flight - score offset. The table in this section contains summary statistics of visual scores by year. The visual score labels and score dates for each plot-level image are accessible at Supplemental Data S1, visual scores from five raters of 100 plots are available in Supplemental Data S2.

## DATA AVAILABILITY STATEMENT

The code for the ArcGIS Pro toolbox used for generating plot outlines can be accessed at https://github.com/reaustin/arcgis-uav-toolbox.git. The code for the orthomosaicing pipeline in the Agisoft Metashape python API can be accessed at https://github.com/nirwan1265/metashape.git. The code for producing plot-level images from orthomosaics can be accessed at https://github.com/gagelab/uav_tools.git. The raw images and processed plot images can be accessed at <AgDataCommons DOI in process>. The code and visual score data for model training, inference, and residual analysis is available at https://github.com/gagelab/uav4slb.git (in process; to be finalized before publication).

## AUTHOR CONTRIBUTIONS

**Cole Hammett:** Conceptualization; data curation; formal analysis; investigation; methodology; software; validation; visualization; writing – original draft; writing – review and editing. **Katelyn Rumley:** Data curation; investigation; methodology; supervision; writing – review and editing. **Peter Balint-Kurti:** Conceptualization; data curation; funding acquisition; investigation; methodology; project administration; resources; supervision; writing – review and editing. **Joseph Gage:** Conceptualization; data curation; funding acquisition; investigation; methodology; project administration; resources; software; supervision; writing – review and editing.

## DECLARATION OF GENERATIVE AI AND AI-ASSISTED TECHNOLOGIES

During the preparation of this work the authors used generative artificial intelligence technologies intermittently during manuscript preparation, coding support, troubleshooting, and literature organization. All outputs were critically reviewed, edited, and validated by the author who takes full responsibility for the integrity and accuracy of the reported results. This was achieved by rigorous exploratory data analysis, human check-ins during processing, and thorough results rationalization using previous related findings. No AI technology was used as an authoritative source of data interpretation, statistical results, or final conclusions. AI technology was used to augment but not replace the author’s scientific judgement.

Two general purpose large language models were employed during the development of this research. OpenAI ChatGPT Plus plan models were used between June 2024 and May 2026 with the model selector beginning at GPT-4 and updating throughout this period to GPT-4o, GPT-4o mini/GPT-4.1 mini, GPT-4.1, OpenAI o3/o4-mini reasoning models, and GPT-5-series models. The default Anthropic Claude models for the Pro plan were used between January 2025 and May 2026 which include Sonnet versions 3.5, 3.7, 4, 4.5, and 4.6. Claude was accessed via the two interfaces: 1) the <u>claude.ai</u> web application and 2) the Claude Code extension for Visual Studio Code (an agentic coding tool). NotebookLM was a retrieval-augmented generation tool powered by Google Gemini that was used for organization, synthesis and summarization of relevant material. Outputs from this AI technology are grounded exclusively in user-uploaded content with in-line citations to the provided sources. All uploaded sources were peer-reviewed literature and publicly available technical documentation.

## ABBREVIATIONS

AGGD: accumulated growing degree days
AI: artificial intelligence
AUDPC: area under disease progress curve
CCRS: Central Crops Research Station
CNN: convolutional neural network
CV: coefficient of variation
DL: deep learning
DPI: days post-inoculation
ECONet: Environment and Climate Observing Network
HSV: hue - saturation - value
HTP: high throughput phenotyping
MSE: mean squared error
OLS: ordinary least squares
QDR: quantitative disease resistance
QTL: quantitative trait loci
RGB: red - green - blue
RMS: root mean square
RMSE: root mean square error
SD: standard deviation
SLB: southern leaf blight
SZA: solar zenith angle
UAV: unmanned aerial vehicle
ViT: vision transformer.

## REFERENCES

Alladassi, B. M. E., Mu, Q., Wei, J., Miguez, F. E., Price, K., Li, X., & Yu, J. (2026). Persistent and transient QTLs underlying growth trajectory of plant height in sorghum. *Journal of Experimental Botany*, erag062. 10.1093/jxb/erag062

Anderson, K. S., Hansen, C. W., Holmgren, W. F., Jensen, A. R., Mikofski, M. A., & Driesse, A. (2023). pvlib python: 2023 project update. Journal of Open Source Software, 8(92), 5994. 10.21105/joss.05994

Araus, J. L., & Cairns, J. E. (2014). Field high-throughput phenotyping: The new crop breeding frontier. Trends in Plant Science, 19(1), 52–61. 10.1016/j.tplants.2013.09.008

Austin, R. (2022). UAV Toolbox (Version 0.1.0) [Python]. https://github.com/reaustin/arcgis-uav-toolbox (Original work published 2022)

Balint-Kurti, P. J., Zwonitzer, J. C., Wisser, R. J., Carson, M. L., Oropeza-Rosas, M. A., Holland, J. B., & Szalma, S. J. (2007). Precise Mapping of Quantitative Trait Loci for Resistance to Southern Leaf Blight, Caused by Cochliobolus heterostrophus Race O, and Flowering Time Using Advanced Intercross Maize Lines. Genetics, 176(1), 645–657. 10.1534/genetics.106.067892

Balint-Kurti, P., & Pataky, J. (2024). Reconsidering the Lessons Learned from the 1970 Southern Corn Leaf Blight Epidemic. Phytopathology®, 114(9), 2007–2016. 10.1094/PHYTO-03-24-0105-PER

Belcher, A. R., Zwonitzer, J. C., Cruz, J. S., Krakowsky, M. D., Chung, C.-L., Nelson, R., Arellano, C., & Balint-Kurti, P. J. (2012). Analysis of quantitative disease resistance to southern leaf blight and of multiple disease resistance in maize, using near-isogenic lines. Theoretical and Applied Genetics, 124(3), 433–445. 10.1007/s00122-011-1718-1

Betts, A. K., Sisson, A. J., Ahumada, D., Allen, T., Anderson, N., Bergstrom, G. C., Bish, M., Bissonnette, K., Broderick, K., Byamukama, E., Chilvers, M. I., Collins, A., Duffeck, M., Esker, P., Faske, T. R., Friskop, A., Harbach, C., Heiniger, R. W., Isakeit, T., … Zeng, Y. (2025). Corn Yield Loss Estimates Due to Diseases in the United States and Ontario, Canada, from 2020 to 2023. Plant Health Progress. 10.1094/PHP-07-25-0193-RS

Bian, Y., Yang, Q., Balint-Kurti, P. J., Wisser, R. J., & Holland, J. B. (2014). Limits on the reproducibility of marker associations with southern leaf blight resistance in the maize nested association mapping population. BMC Genomics, 15(1), 1068. 10.1186/1471-2164-15-1068

Bock, C. H., Barbedo, J. G. A., Del Ponte, E. M., Bohnenkamp, D., & Mahlein, A.-K. (2020). From visual estimates to fully automated sensor-based measurements of plant disease severity: Status and challenges for improving accuracy. Phytopathology Research, 2(1), 9. 10.1186/s42483-020-00049-8

Bock, C. H., Chiang, K.-S., & Del Ponte, E. M. (2022). Plant disease severity estimated visually: A century of research, best practices, and opportunities for improving methods and practices to maximize accuracy. Tropical Plant Pathology, 47(1), 25–42. 10.1007/s40858-021-00439-z

Bock, C. H., Parker, P. E., Cook, A. Z., & Gottwald, T. R. (2008). Characteristics of the Perception of Different Severity Measures of Citrus Canker and the Relationships Between the Various Symptom Types. Plant Disease, 92(6), 927–939. 10.1094/PDIS-92-6-0927

Bongomin, O., Lamo, J., Guina, J. M., Okello, C., Ocen, G. G., Obura, M., Alibu, S., Owino, C. A., Akwero, A., & Ojok, S. (2024). UAV image acquisition and processing for high-throughput phenotyping in agricultural research and breeding programs. The Plant Phenome Journal, 7(1), e20096. 10.1002/ppj2.20096

*Cardinal* [data retrieval interface]. (n.d.). [Dataset]. Retrieved March 9, 2026, from https://products.climate.ncsu.edu/cardinal/request

Carson, M. L. (1998). Aggressiveness and Perennation of Isolates of Cochliobolus heterostrophus from North Carolina. Plant Disease, 82(9), 1043–1047. 10.1094/PDIS.1998.82.9.1043

Carson, M. L., Stuber, C. W., & Senior, M. L. (2004). Identification and Mapping of Quantitative Trait Loci Conditioning Resistance to Southern Leaf Blight of Maize Caused by Cochliobolus heterostrophus Race O. Phytopathology®, 94(8), 862–867. 10.1094/PHYTO.2004.94.8.862

Chakhvashvili, E., Machwitz, M., Antala, M., Rozenstein, O., Prikaziuk, E., Schlerf, M., Naethe, P., Wan, Q., Komárek, J., Klouek, T., Wieneke, S., Siegmann, B., Kefauver, S., Kycko, M., Balde, H., Paz, V. S., Jimenez-Berni, J. A., Buddenbaum, H., Hänchen, L., … Rascher, U. (2024). Crop stress detection from UAVs: Best practices and lessons learned for exploiting sensor synergies. Precision Agriculture, 25(5), 2614–2642. 10.1007/s11119-024-10168-3

Chen, G., Xiao, Y., Dai, S., Dai, Z., Wang, X., Li, B., Jaqueth, J. S., Li, W., Lai, Z., Ding, J., & Yan, J. (2023). Genetic basis of resistance to southern corn leaf blight in the maize multi parent population and diversity panel. Plant Biotechnology Journal, 21(3), 506–520. 10.1111/pbi.13967

Craze, H. A., Pillay, N., Joubert, F., & Berger, D. K. (2022). Deep Learning Diagnostics of Gray Leaf Spot in Maize under Mixed Disease Field Conditions. Plants, 11(15). 10.3390/plants11151942

Dai, Z., Liu, H., Le, Q. V., & Tan, M. (2021). *CoAtNet: Marrying Convolution and Attention for All Data Sizes* (arXiv:2106.04803). arXiv. 10.48550/arXiv.2106.04803

DeChant, C., Wiesner-Hanks, T., Chen, S., Stewart, E. L., Yosinski, J., Gore, M. A., Nelson, R. J., & Lipson, H. (2017). Automated Identification of Northern Leaf Blight-Infected Maize Plants from Field Imagery Using Deep Learning. Phytopathology®, 107(11), 1426–1432. 10.1094/PHYTO-11-16-0417-R

Erenstein, O., Jaleta, M., Sonder, K., Mottaleb, K., & Prasanna, B. M. (2022). Global maize production, consumption and trade: Trends and R&D implications. Food Security, 14(5), 1295–1319. 10.1007/s12571-022-01288-7

Fang, Y., Sun, Q., Wang, X., Huang, T., Wang, X., & Cao, Y. (2024a). EVA-02: A Visual Representation for Neon Genesis. Image and Vision Computing, 149, 105171. 10.1016/j.imavis.2024.105171

Fang, Y., Sun, Q., Wang, X., Huang, T., Wang, X., & Cao, Y. (2024b). EVA-02: A Visual Representation for Neon Genesis. Image and Vision Computing, 149, 105171. 10.1016/j.imavis.2024.105171

Gage, J. L., de Leon, N., & Clayton, M. K. (2018). Comparing Genome-Wide Association Study Results from Different Measurements of an Underlying Phenotype. G3 Genes|Genomes|Genetics, 8(11), 3715–3722. 10.1534/g3.118.200700

Green, J. M., Appel, H., Rehrig, E. M., Harnsomburana, J., Chang, J.-F., Balint-Kurti, P., & Shyu, C.-R. (2012). PhenoPhyte: A flexible affordable method to quantify 2D phenotypes from imagery. Plant Methods, 8(1), 45. 10.1186/1746-4811-8-45

Guo, X., Li, Q., Morrison-Smith, S., Anthony, L., Zare, A., & Song, Y. (2024). Elicitating Challenges and User Needs Associated with Annotation Software for Plant Phenotyping. Proceedings of the 29th International Conference on Intelligent User Interfaces, IUI ’24, 431–443. 10.1145/3640543.3645178

Haque, M. A., Marwaha, S., Arora, A., Paul, R. K., Hooda, K. S., Sharma, A., & Grover, M. (2021). Image-based identification of maydis leaf blight disease of maize (Zea mays) using deep learning. The Indian Journal of Agricultural Sciences, 91(9), 1362–1367. 10.56093/ijas.v91i9.116089

Holley, R. N. (1989). New Sources of Resistance to Southern Corn Leaf Blight from Tropical Hybrid Maize Derivatives. Plant Disease, 73(7), 562. 10.1094/PD-73-0562

Hu, X., Chen, S., Duan, Q., Ahn, C. K., Shang, H., & Zhang, D. (2025). *Domain Adaptation for Big Data in Agricultural Image Analysis: A Comprehensive Review* (arXiv:2506.05972). arXiv. 10.48550/arXiv.2506.05972

Ispizua Yamati, F. R., Günder, M., Bömer, J., Barreto, A., Varrelmann, M., Bauckhage, C., & Mahlein, A.-K. (2026). Hybrid Modeling of Cercospora Leaf Spot Epidemiology: Integrating Mechanistic and Machine Learning Approaches Using Remote-Sensing and Environmental Data. Phytopathology®, 116(4), 561–578. 10.1094/PHYTO-03-25-0113-R

Jarquín, D., Lemes da Silva, C., Gaynor, R. C., Poland, J., Fritz, A., Howard, R., Battenfield, S., & Crossa, J. (2017). Increasing Genomic-Enabled Prediction Accuracy by Modeling Genotype × Environment Interactions in Kansas Wheat. The Plant Genome, 10(2), plantgenome2016.12.0130. 10.3835/plantgenome2016.12.0130

Jeger, M., Beresford, R., Berlin, A., Bock, C., Fox, A., Gold, K. M., Newton, A. C., Vicent, A., & Xu, X. (2024). Impact of novel methods and research approaches in plant pathology: Are individual advances sufficient to meet the wider challenges of disease management? Plant Pathology, 73(7), 1629–1655. 10.1111/ppa.13927

Kump, K. L., Bradbury, P. J., Wisser, R. J., Buckler, E. S., Belcher, A. R., Oropeza-Rosas, M. A., Zwonitzer, J. C., Kresovich, S., McMullen, M. D., Ware, D., Balint-Kurti, P. J., & Holland, J. B. (2011). Genome-wide association study of quantitative resistance to southern leaf blight in the maize nested association mapping population. Nature Genetics, 43(2), 163–168. Agricultural & Environmental Science Collection; ProQuest Central; SciTech Premium Collection (849563456; 21217757).

Large, E. C. (1966). Measuring Plant Disease. Annual Review of Phytopathology, 4(Volume 4, 1966), 9–26. 10.1146/annurev.py.04.090166.000301

Lee, D.-Y., Na, D.-Y., Góngora-Canul, C., Jimenez-Beitia, F. E., Goodwin, S. B., Cruz, A. P., Delp, E. J., Acosta, A. G., Lee, J.-S., Falconí, C. E., & Cruz, C. D. (2025). Optimizing Corn Tar Spot Measurement: A Deep Learning Approach Using Red-Green-Blue Imaging and the Stromata Contour Detection Algorithm for Leaf-Level Disease Severity Analysis. Plant Disease, 109(1), 73–83. 10.1094/PDIS-12-23-2702-RE

Li, J., Wu, W., Zhao, C., Bai, X., Dong, L., Tan, Y., Yusup, M., Akelebai, G., Dong, H., & Zhi, J. (2025). Effects of solar elevation angle on the visible light vegetation index of a cotton field when extracted from the UAV. Scientific Reports, 15(1), 18497. 10.1038/s41598-025-00992-6

Li, Y., Chen, L., Li, C., Bradbury, P. J., Shi, Y., Song, Y., Zhang, D., Zhang, Z., Buckler, E. S., Li, Y., & Wang, T. (2018). Increased experimental conditions and marker densities identified more genetic loci associated with southern and northern leaf blight resistance in maize. Scientific Reports, 8(1), 6848. 10.1038/s41598-018-25304-z

Lim, S., & Hooker, A. (1976). Estimates of combining ability for resistance to Helminthosporium maydis race 0 in a maize population. Maydica, 21(3), 121–128. CABI Databases.

Liu, Z., Hu, H., Lin, Y., Yao, Z., Xie, Z., Wei, Y., Ning, J., Cao, Y., Zhang, Z., Dong, L., Wei, F., & Guo, B. (2022). *Swin Transformer V2: Scaling Up Capacity and Resolution* (arXiv:2111.09883). arXiv. 10.48550/arXiv.2111.09883

Madden, L. V., Hughes, G., & van den Bosch, F. (2017). CHAPTER 2: Measuring Plant Diseases. In H. Gareth, van den B. Frank, & V. M. Laurence (Eds.), The Study of Plant Disease Epidemics (pp. 11–31). The American Phytopathological Society. 10.1094/9780890545058.002

Mardanisamani, S., & Eramian, M. (2022). Segmentation of vegetation and microplots in aerial agriculture images: A survey. The Plant Phenome Journal, 5(1), e20042. 10.1002/ppj2.20042

Martins, L. B., Rucker, E., Thomason, W., Wisser, R. J., Holland, J. B., & Balint-Kurti, P. (2019). Validation and Characterization of Maize Multiple Disease Resistance QTL. G3 Genes|Genomes|Genetics, 9(9), 2905–2912. 10.1534/g3.119.400195

Munkvold, G. P., & White, D. G. (2016). PART I: Infectious Diseases. In Compendium of Corn Diseases, Fourth Edition (pp. 7–130). The American Phytopathological Society. 10.1094/9780890544945.002

Negeri, A. T., Coles, N. D., Holland, J. B., & Balint-Kurti, P. J. (2011). Mapping QTL Controlling Southern Leaf Blight Resistance by Joint Analysis of Three Related Recombinant Inbred Line Populations. Crop Science, 51(4), 1571–1579. 10.2135/cropsci2010.12.0672

Nelson, R., Wiesner-Hanks, T., Wisser, R., & Balint-Kurti, P. (2018). Navigating complexity to breed disease-resistant crops. Nature Reviews Genetics, 19(1), 21–33. 10.1038/nrg.2017.82

Oquab, M., Darcet, T., Moutakanni, T., Vo, H., Szafraniec, M., Khalidov, V., Fernandez, P., Haziza, D., Massa, F., El-Nouby, A., Assran, M., Ballas, N., Galuba, W., Howes, R., Huang, P.-Y., Li, S.-W., Misra, I., Rabbat, M., Sharma, V., … Bojanowski, P. (2024). *DINOv2: Learning Robust Visual Features without Supervision* (arXiv:2304.07193). arXiv. 10.48550/arXiv.2304.07193

Poland, J. A., Balint-Kurti, P. J., Wisser, R. J., Pratt, R. C., & Nelson, R. J. (2009). Shades of gray: The world of quantitative disease resistance. Trends in Plant Science, 14(1), 21–29. 10.1016/j.tplants.2008.10.006

Poland, J. A., & Nelson, R. J. (2011). In the Eye of the Beholder: The Effect of Rater Variability and Different Rating Scales on QTL Mapping. Phytopathology®, 101(2), 290–298. 10.1094/PHYTO-03-10-0087

Rife, T. W., & Poland, J. A. (2014). Field Book: An Open-Source Application for Field Data Collection on Android. Crop Science, 54(4), 1624–1627. 10.2135/cropsci2013.08.0579

Schwanck, A. A., & Del Ponte, E. M. (2014). Accuracy and Reliability of Severity Estimates Using Linear or Logarithmic Disease Diagram Sets in True Colour or Black and White: A Study Case for Rice Brown Spot. Journal of Phytopathology, 162(10), 670–682. 10.1111/jph.12246

Sermons, S. M., & Balint-Kurti, P. J. (2018). Large Scale Field Inoculation and Scoring of Maize Southern LeafBlight and Other Maize Foliar Fungal Diseases. Bio-Protocol, 8(5), e2745. 10.21769/BioProtoc.2745

Shahi, T. B., Xu, C.-Y., Neupane, A., & Guo, W. (2023). Recent Advances in Crop Disease Detection Using UAV and Deep Learning Techniques. Remote Sensing, 15(9). 10.3390/rs15092450

Shahi, T. B., Xu, C.-Y., Neupane, A., Guo, W., Shahi, T. B., Xu, C.-Y., Neupane, A., & Guo, W. (2022). Machine learning methods for precision agriculture with UAV imagery: A review. Electronic Research Archive, 30(12), 4277–4317. 10.3934/era.2022218

Sherwood, R. T. (1983). Illusions in Visual Assessment of Stagonospora Leaf Spot of Orchardgrass. Phytopathology, 73(2), 173. 10.1094/Phyto-73-173

Singh, A., Ganapathysubramanian, B., Singh, A. K., & Sarkar, S. (2016). Machine Learning for High-Throughput Stress Phenotyping in Plants. Trends in Plant Science, 21(2), 110–124. 10.1016/j.tplants.2015.10.015

Singh, R. K., Tiwari, A., & Gupta, R. K. (2022). Deep transfer modeling for classification of Maize Plant Leaf Disease. Multimedia Tools and Applications, 81(5), 6051–6067. 10.1007/s11042-021-11763-6

Spisak, T. (2022). Statistical quantification of confounding bias in machine learning models. GigaScience, 11, giac082. 10.1093/gigascience/giac082

Stewart, E. L., Wiesner-Hanks, T., Kaczmar, N., DeChant, C., Wu, H., Lipson, H., Nelson, R. J., & Gore, M. A. (2019). Quantitative Phenotyping of Northern Leaf Blight in UAV Images Using Deep Learning. Remote Sensing, 11(19). 10.3390/rs11192209

Tan, M., & Le, Q. V. (2021). *EfficientNetV2: Smaller Models and Faster Training* (arXiv:2104.00298). arXiv. 10.48550/arXiv.2104.00298

Tu, Z., Talebi, H., Zhang, H., Yang, F., Milanfar, P., Bovik, A., & Li, Y. (2022). *MaxViT: Multi-Axis Vision Transformer* (arXiv:2204.01697). arXiv. 10.48550/arXiv.2204.01697

Ullstrup, A. J. (1972). The Impacts of the Southern Corn Leaf Blight Epidemics of 1970-1971. Annual Review of Phytopathology, 10(Volume 10,), 37–50. 10.1146/annurev.py.10.090172.000345

Valencia-Ortiz, M., Sangjan, W., Selvaraj, M. G., McGee, R. J., & Sankaran, S. (2021). Effect of the Solar Zenith Angles at Different Latitudes on Estimated Crop Vegetation Indices. Drones, 5(3). 10.3390/drones5030080

Wang, X., Choi, D., Adams, D., Ahn, J., Balmant, K., Dias, R., Messina, C. D., Muñoz-Carpena, R., Whitaker, V. M., Yu, H., Yu, Z., Zhao, C., & Li, C. (2026). Artificial intelligence-powered plant phenomics: Progress, challenges, and opportunities. The Plant Phenome Journal, 9(1), e70060. 10.1002/ppj2.70060

Wiesner-Hanks, T., Wu, H., Stewart, E., DeChant, C., Kaczmar, N., Lipson, H., Gore, M. A., & Nelson, R. J. (2019). Millimeter-Level Plant Disease Detection From Aerial Photographs via Deep Learning and Crowdsourced Data. Frontiers in Plant Science, Volume 10-2019. 10.3389/fpls.2019.01550

Wisser, R. J., Balint-Kurti, P. J., & Nelson, R. J. (2006). The Genetic Architecture of Disease Resistance in Maize: A Synthesis of Published Studies. Phytopathology®, 96(2), 120– 129. 10.1094/PHYTO-96-0120

Woo, S., Debnath, S., Hu, R., Chen, X., Liu, Z., Kweon, I. S., & Xie, S. (2023). *ConvNeXt V2: Co-designing and Scaling ConvNets with Masked Autoencoders* (arXiv:2301.00808). arXiv. 10.48550/arXiv.2301.00808

Wu, H., Wiesner-Hanks, T., Stewart, E. L., DeChant, C., Kaczmar, N., Gore, M. A., Nelson, R. J., & Lipson, H. (2019). Autonomous Detection of Plant Disease Symptoms Directly from Aerial Imagery. The Plant Phenome Journal, 2(1), 190006. 10.2135/tppj2019.03.0006

Yang, Q., Balint-Kurti, P., & Xu, M. (2017). Quantitative Disease Resistance: Dissection and Adoption in Maize. Molecular Plant, 10(3), 402–413. 10.1016/j.molp.2017.02.004

Yang, Q., He, Y., Kabahuma, M., Chaya, T., Kelly, A., Borrego, E., Bian, Y., El Kasmi, F., Yang, L., Teixeira, P., Kolkman, J., Nelson, R., Kolomiets, M., L Dangl, J., Wisser, R., Caplan, J., Li, X., Lauter, N., & Balint-Kurti, P. (2017). A gene encoding maize caffeoyl-CoA O-methyltransferase confers quantitative resistance to multiple pathogens. Nature Genetics, 49(9), 1364–1372. 10.1038/ng.3919

Yang, W., Feng, H., Zhang, X., Zhang, J., Doonan, J. H., Batchelor, W. D., Xiong, L., & Yan, J. (2020). Crop Phenomics and High-Throughput Phenotyping: Past Decades, Current Challenges, and Future Perspectives. Molecular Plant, 13(2), 187–214. 10.1016/j.molp.2020.01.008

Yin, C., Zeng, T., Zhang, H., Fu, W., Wang, L., & Yao, S. (2022). Maize Small Leaf Spot Classification Based on Improved Deep Convolutional Neural Networks with a Multi-Scale Attention Mechanism. Agronomy, 12(4). 10.3390/agronomy12040906

Zhu, H., Huang, Y., An, Z., Zhang, H., Han, Y., Zhao, Z., Li, F., Zhang, C., & Hou, C. (2024). Assessing radiometric calibration methods for multispectral UAV imagery and the influence of illumination, flight altitude and flight time on reflectance, vegetation index and inversion of winter wheat AGB and LAI. Computers and Electronics in Agriculture, 219, 108821. 10.1016/j.compag.2024.108821

Zwonitzer, J. C., Bubeck, D. M., Bhattramakki, D., Goodman, M. M., Arellano, C., & Balint-Kurti, P. J. (2009). Use of selection with recurrent backcrossing and QTL mapping to identify loci contributing to southern leaf blight resistance in a highly resistant maize line. Theoretical and Applied Genetics, 118(5), 911–925. 10.1007/s00122-008-0949-2

